# Measuring and Mitigating PCR Bias in Microbiome Data

**DOI:** 10.1101/604025

**Authors:** Justin D. Silverman, Rachael J. Bloom, Sharon Jiang, Heather K. Durand, Sayan Mukherjee, Lawrence A. David

## Abstract

PCR amplification plays a central role in the measurement of mixed microbial communities via high-throughput sequencing. Yet PCR is also known to be a common source of bias in microbiome data. Here we present a paired modeling and experimental approach to characterize and mitigate PCR bias in microbiome studies. We use experimental data from mock bacterial communities to validate our approach and human gut microbiota samples to characterize PCR bias under real-world conditions. Our results suggest that PCR can bias estimates of microbial relative abundances by a factor of 2-4 but that this bias can be mitigated using simple Bayesian multinomial logistic-normal linear models.

**Author summary:** High-throughput sequencing is often used to profile host-associated microbial communities. Many processing steps are required to transform a community of bacteria into a pool of DNA suitable for sequencing. One important step is amplification where, to create enough DNA for sequencing, DNA from many different bacteria are repeatedly copied using a technique called Polymerase Chain Reaction (PCR). However, PCR is known to introduce bias as DNA from some bacteria are more efficiently copied than others. Here we introduce an experimental procedure that allows this bias to be measured and computational techniques that allow this bias to be mitigated in sequencing data.

## Introduction

Polymerase Chain Reaction (PCR) amplification is an essential procedure used when profiling microbial communities by high-throughput sequencing. Yet, PCR is also known to be a common source of bias in high-throughput sequencing studies [1–3]. In the study of microbial communities by 16S rRNA, PCR bias can be substantial. Mock communities have been used to demonstrate that over-amplification of specific 16S rRNA templates occurs reproducibly, often with preferential amplification of over 3.5 fold [4]. Even single nucleotide mismatches between primer and template have been shown to lead to preferential amplification of up to 10 fold [5].

Relatively little progress though has been made in characterizing PCR bias in such a way that the bias can be corrected. Dual-indexing amplification with high-fidelity polymerases has been shown to lead to decreased PCR bias in microbiome studies [6]. Such experimental modifications can still have high levels of error [6]. Recently, a computational method called *alpine* was proposed as a means of inferring and correcting PCR bias in RNA-seq studies [3]. However, *alpine* requires the presence of reference genomes against which transcripts can be aligned, something that is often not present in the analysis of metagenomic or amplicon based microbiome studies which characterize multiple taxa simultaneously.

Here we introduce a calibration curve for PCR which allows bias to be characterized directly from host associated microbial communities without the need to create mock standards. We pair with this calibration curve with Bayesian multinomial logistic-normal linear (*pibble*) models which learn and mitigate PCR bias while accounting for uncertainty due to multivariate counting and random technical variation [7,8]. We validate our approach using both mock and human gut microbial communities. Our results support the hypotheses that substantial bias is introduced when using DNA amplification to survey microbial communities, as well as demonstrate how a simple modeling method can help mitigate this bias.

## Results

### Measuring and Modeling PCR Bias

To develop a model of PCR bias we denote by *α_j_* the absolute abundance of a transcript *j* ∈ {1, …, *D*} in a pool of DNA prior to PCR amplification. We also denote by *b_j_* the efficiency with which transcript *j* is amplified by PCR, *e.g*., *b_j_* = 2 implies that transcript *j* undergoes perfect doubling at each PCR cycle. Finally, we denote by *w_ij_* the absolute abundance of a transcript *j* in a pool of DNA after *x_i_* cycles of PCR. With this notation we can write the following multiplicative model for PCR bias:

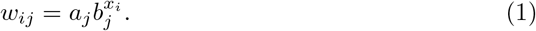

Equation (1) can be written in log scale using vector notation as the following log-linear model:

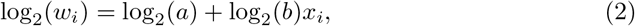

where log_2_(*w_i_*) refers to the element-wise logarithm of the vector *w_i_*. Equation (2) states that bias (deviation from doubling at each cycle) would present as a non-zero slope of a regression line relating microbial abundance to PCR cycle number (log_2_(*b*) ≠ 0).

This model suggests that, given measurements of transcript abundance (*w_i_*) at different PCR cycle numbers (*x_i_*), we could infer the unbiased abundance of each transcript (a) and the efficiency or bias with which each transcript is amplified (b). We therefore propose creating calibration curves that utilize multiple aliquots of DNA extracted from the same microbial community, which are then amplified with varying numbers of PCR cycles and sequenced using high-throughput multiplex sequencing (HTS).

### Mock Community Analysis

To evaluate the utility of our approach to characterizing and removing PCR bias, we designed a mock community where DNA from fourteen bacterial isolates was pooled in approximately known amounts. To capture PCR bias, the mock community was split into aliquots and each aliquot underwent a predetermined number of PCR cycles varying from 3 to 35 cycles. To avoid systematic bias from the ordering in which the amplifications were done, the order of PCRs were randomized. The resulting amplified DNA was pooled and sequenced. We found that we could only reliably map five 16S rRNA sequences in our HTS pipeline to mock community members; reads from isolates that could not be uniquely mapped were amalgamated into a category called “other”.

The resulting table of sequence variants was analyzed using a multinomial logistic-normal linear (*pibble*) model (*Methods*). This model accounts for the composition nature of 16S rRNA HTS data [9,10] as well as uncertainty due to multivariate counting [8]. We also added to our model a binary covariate denoting whether samples were amplified in the first or second batch of PCR reactions to account for sample processing batch effects. Finally, based on prior reports of the accuracy of qPCR [11], we made the assumption that our measurement of the true microbial composition of this mock community was accurate to within one order of magnitude in log-ratio space.

As indicated by Equation (2), a non-zero slope in the relationship between community composition and PCR cycle number would be evidence for PCR bias. Visual inspection confirmed a linear relationship between these variables according to both the posterior marginal (Figure 1) and the posterior predictive distribution (Figure S1) of the fitted *pibble* model. Moreover, the slope of this relationship indicates that indeed, bias was introduced over 35 cycels of PCR (Figure 1) – relative bias approached 4-8 fold for some sequence variants (Figure S2).

**Fig 1.**
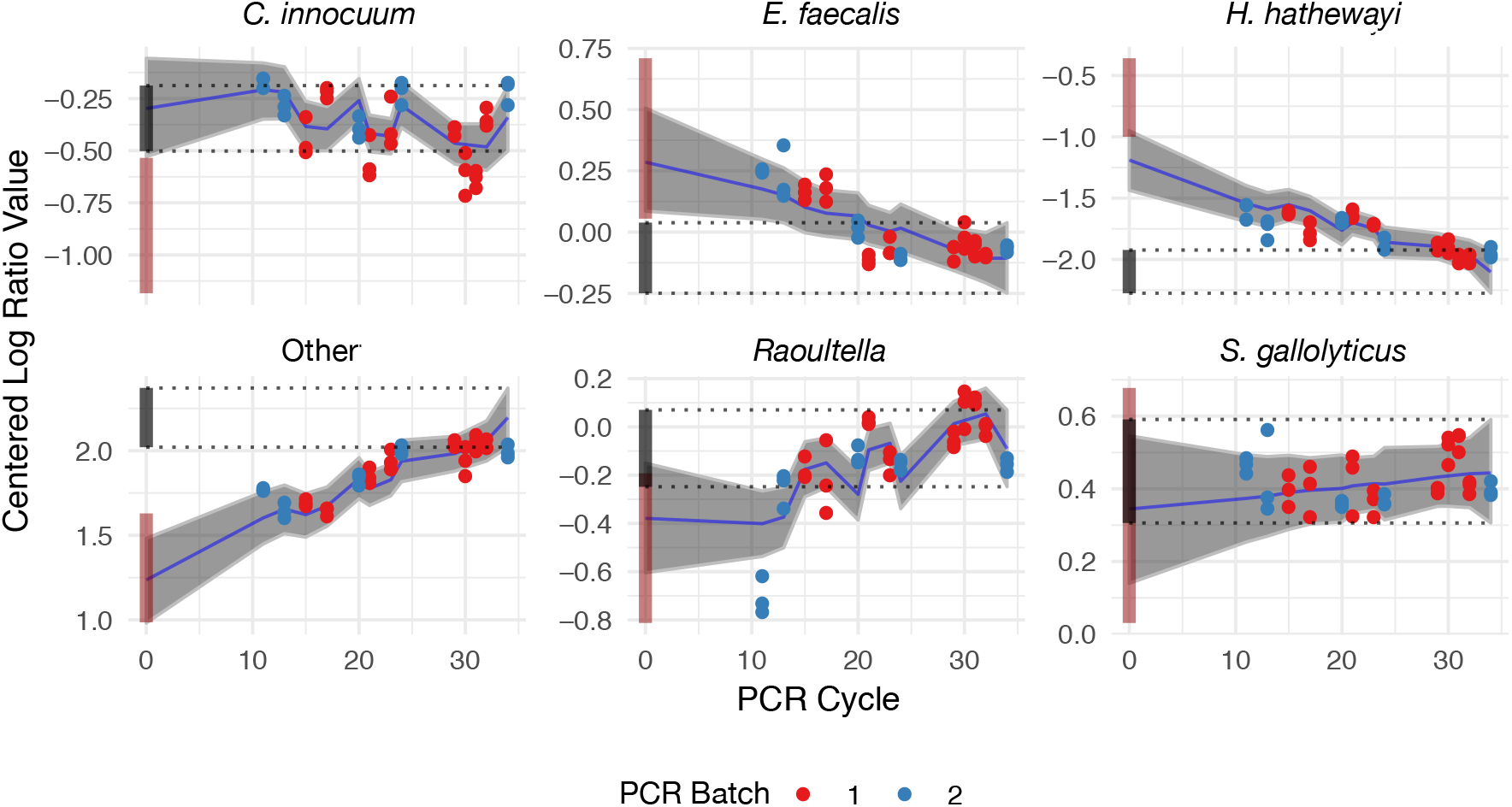
Combining calibration experiments with multinomial logistic-normal linear (*pibble*) models allows PCR bias to be mitigated. Mean (blue line) and 95% credible regions (grey ribbon) from *pibble* model (Λ*X*; *Methods*) applied to mock community calibration data. To illustrate the impact of PCR bias the compositional estimates from cycle 35 were projected onto cycle 0 (cycle 0 is the unamplified community; black bar). The inferred linear relationship between PCR cycle number and composition is shown after adjusting for PCR batch. This adjustment leads the linear fit to appear non-linear even though the overall model is linear. A pseudo-count of 0.65 was added to observed count data prior to log-ratio transformation to enable the data to be visualized along with the posterior estimates. This pseudo-count was included for visualization purposes only and was not required for fitting the *pibble* model. The true mock community composition is shown with measurement error as a dark red bar at PCR cycle 0.

Still, by using our *pibble* model fit to this calibration curve data, we found that we could remove much of the bias introduced by PCR (Figure 1). Our model removed bias in 4 of 6 log-ratios and correctly inferred that one log-ratio had little bias (*S. gallolyticus*). Only one log-ratio, the coordinate corresponding to *C. innocuum*, remained uncorrected by our model. However, we note that the overall bias of *C. innocuum* was slight and that our model did not worsen the bias. Thus, in 6 out of 6 log-ratios, our model either left the bias unchanged or mitigated it.

### Human Gut Microbial Community Analysis

To characterize and correct PCR bias in human gut microbial communities we repeated the experimental approach used for the mock community but applied to four different communities derived from human hosts. Each community was cultured *ex vivo* for 1-3 days using an independent artificial gut systems as previously described [8]. The PCR experiments for these real communities were performed on multiple PCR machines due to the large number of samples involved. After initial preprocessing, the resulting data represented 68 bacterial genera from 6 bacterial phyla. To fit this data, we modeled each of the four individuals with random intercepts, a fixed effect for cycle number, and random effects for each PCR machine (*Methods*).

As in our analysis of the mock community, we find that the calibration data from human gut microbial communities is well fit by a *pibble* model (Figure S3 and File S1). This further supports our conceptual model for PCR bias in human gut microbial community data. To succinctly visualize the scale of PCR bias present when amplifying human gut microbial communities, we investigated the total bias introduced into the data after 35 cycles of PCR (Figure 2). As in our evaluation of the mock community, we find that 35 cycles of PCR induces a substantial bias in estimated relative abundances with approximately 15% of taxa being subject to over a factor of 2 bias (Figure S4). These results are in line with prior reports suggesting an average deviation between 2-3 fold [6]. Our results suggest that the genera *Holdemania, Coprococcus*, and *Ruminococcus* are consistently the most under-represented taxa due to PCR bias while *Parasutterella, Bifidobacterium*, and *Bacteroides* are consistently the most over-represented.

**Fig 2.**
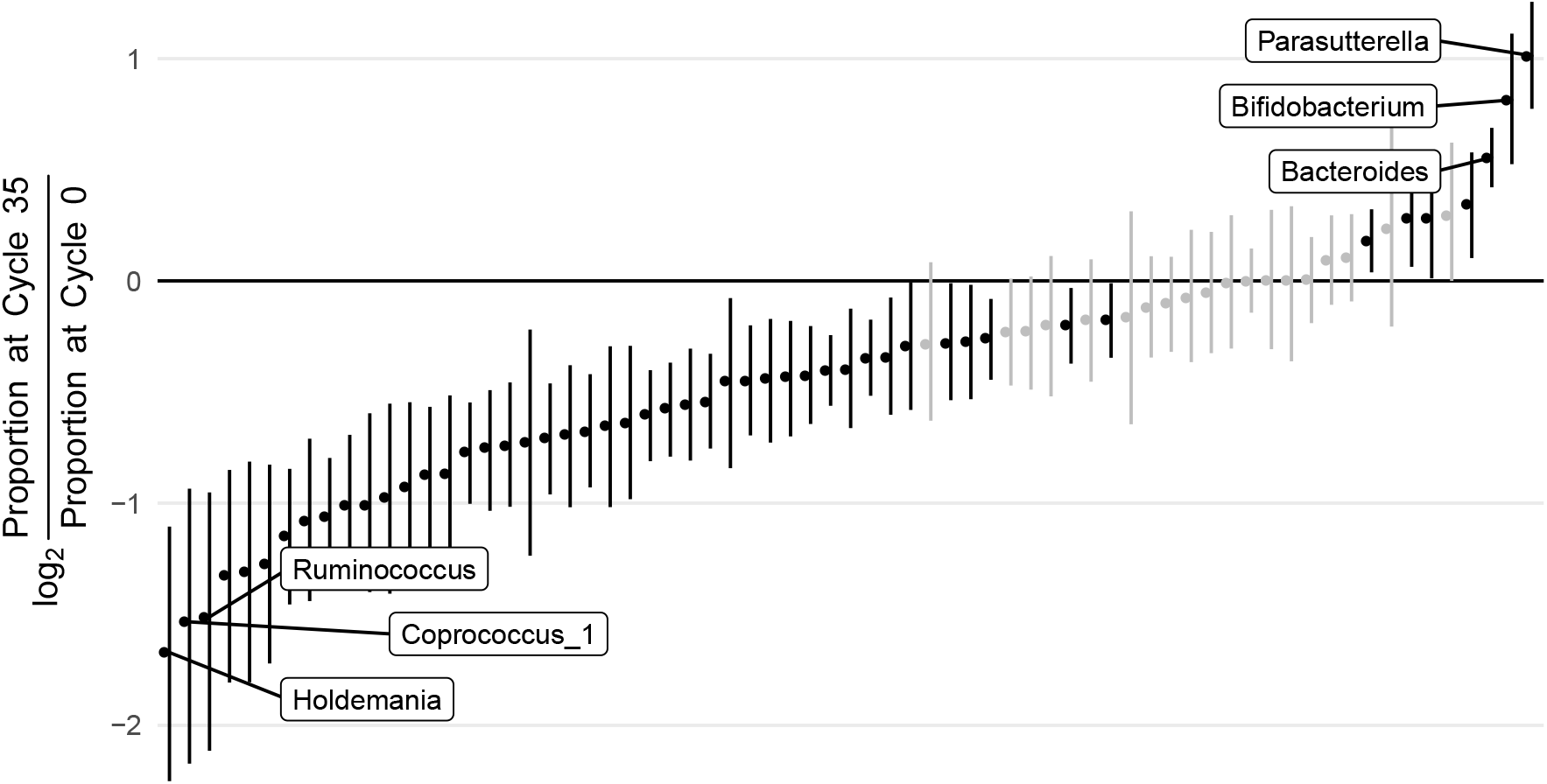
Evidence that PCR induces substantial bias in human gut microbial communities. To visualize the scale of PCR bias in real microbial communities we calculated bias induced after 35 cycles of PCR as the log-ratio of the taxon proportion at cycle 35 versus inferred taxon proportions at cycle 0 (unamplified). The mean and 95% credible regions for this bias is depicted for each taxon. Those taxa with 95% credible regions not overlapping zero are shown in black. This absolute value of this bias on the proportional scale is presented in Figure S4. A full set of posterior fits similar to those shown in Figure 1 is given in File S1.

Investigating modelled random effects associated with PCR machine revealed that PCR reactions run on one machine were substantially different than those run on the other machines (Figure S5). Reviewing the settings on each PCR machine, it was found that the outlying machine had its temperature mis-set during the annealing phase of each PCR cycle *(Methods*). Our bias estimates therefore excluded samples from this PCR machine. More broadly, this finding demonstrates how creating PCR calibration curves can be used to detect and correct for sample processing errors in microbiota surveys.

Last, we hypothesized that PCR bias could be predicted based on either sequence similarity (using the Levenshtein distance) or GC content of 16S rRNA amplicons. While the primer binding sequence has been associated with PCR bias [4,12], this sequence is often not available from standard bioinformatics pipelines whereas the 16S rRNA amplicon sequence is. Therefore, if such prediction was possible, it could provide a means of mitigating PCR bias for taxa not included in an initial calibration experiment with minimal additional effort. To investigate this hypothesis we used both linear and non-linear models to predict bias based on these metrics *(Methods*). Both sequence similarity and GC content of inferred amplicons proved to be poor predictors of PCR bias (see *Methods* for further details). This failure of prediction suggests that PCR bias is more complicated than GC content or sequence similarity of sequenced amplicons and may require access to the primer binding sequence for prediction.

## Discussion

Here we have presented an approach to characterizing and correcting PCR bias in microbiome studies based on a simple calibration experiment and multinomial logistic-normal linear (*pibble*) models. Using both mock and human gut microbial communities we demonstrated that sequencing data were well-fit by log-linear models, which lends credence to our conceptual model of PCR bias as a multiplicative process. Moreover, our model suggests that PCR can bias relative abundance estimates by a factor of 2-8. Still, using mock communities we demonstrated that our approach can measure and mitigate PCR bias. Although we used these mock communities to validate our approach, our approach does not require that mock communities be used to measure and mitigate bias in microbiota survey data. We find this appealing as many microbial taxa that may be of interest are difficult to isolate and culture without specialized experimental techniques [13].

While our mock community results demonstrated we can mitigate PCR bias it also suggested that our current methods may not completely remove this bias. In particular there were two log-ratios, the log-ratio coordinates for *C. innocuum* and *H. hathewayi* that may have be under-corrected. There are two potential explanations for this. First, other sources of bias and random variation which we have not accounted for may be contributing to the observed data [14]. For example, bias present in DNA sequencing, which also relies on amplification [15], could affect measurements but would not have been captured by our calibration experiments. Second, our conceptual model of PCR bias as a multiplicative process may fail to account for some subtleties of this error. For example, it has been demonstrated that template annealing of high abundance sequences in the later stages of amplification could inhibit application [1,16]. We would expect such abundance dependent effects to appear non-linear in log-ratio space. Future studies investigating other sources of technical bias and non-log-linear aspects of PCR bias would likely provide avenues for more completely removing PCR bias.

Even without further refinement, we believe that this work provides a simple experimental and computational approach for mitigating PCR bias in real data without the need for mock community standards. For those wishing to apply this method we recommend that this calibration experiment be performed using a pooled library of samples. Such pooling would ensure that the bias associated with each taxon in a study can be captured by the calibration experiment. Additionally, we recommend the collection of technical replicates within the calibration experiment as demonstrated in both our mock and human gut microbial community experiments. As PCR bias is just one source of technical variation [14], technical replicates may prevent other sources of technical variation from being incorrectly attributed to PCR bias.

## Materials and methods

### PCR Bias Model

To account for the fact that high throughput-sequencing reflects the relative abundance of microbial taxa in a community [9,10], we constrain the model in Equation (2) to the simplex – the mathematical space describing relative abundances. This constraint can be imposed by multiplying Equation (2) by a *D* − 1 × *D* contrast matrix Ψ to so that Equation (2) is parameterized by log-ratios:

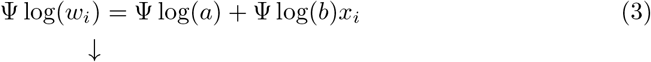

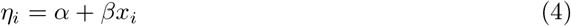

where *η* now represents the relative abundance corresponding to *w_i_* but represented as a vector of log-ratios determined by the contrast matrix Ψ.

Beyond PCR bias, sequence count data may be subject to other sources of technical variation including but not limited to variation from counting [17] and batch effects. To account for these sources of random variation we embed Equation (4) in the following probabilistic model

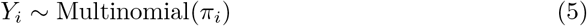

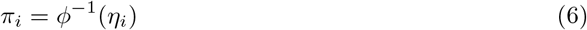

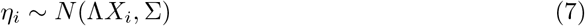

where *Y_i_* denotes the sequence counts from a sample *i* ∈ {1, …, *D*}, Λ*X_i_* denotes a generalization of *α* + *βx_i_* to a larger class of linear models (*e.g*., allowing other covariates such as batch number to be modeled in addition to PCR cycle number), and *ϕ*^−1^(*η_i_*) denotes the inverse transformation of *η_i_* = Ψ log(*π_i_*) which is given by 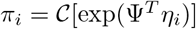 and where 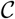 denotes the closure operation defined as

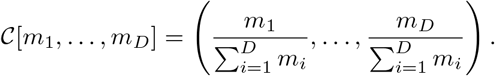

Equations (5)–(7) denote a multinomial logistic-normal linear model similar to that proposed by Silverman et al. [8] as part of the MALLARD framework for time-series analysis of microbiome data. In this work we fit a Bayesian formulation of the above model using matrix-normal and inverse Wishart priors

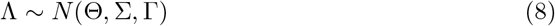

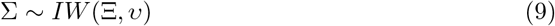

which is available as the function *pibble* in the *stray* R package [18] which uses a marginal Laplace approximation for inference [7]. Together, Equations (5)–(9) form a generative model for PCR bias in sequence count data motivated by the multiplicative model of PCR bias given in Equation (2).

### Sample Acquisition

Fecal samples were collected from four human subjects under a protocol approved by the Duke Health Institutional Review Board (Duke Health IRB Pro00049498). Subjects provided fecal samples at no risk to themselves, had no acute enteric illness, and had not taken antibiotics in the past month.

### Mock Community Data Collection

The mock community included DNA from fourteen isolates; six were obtained from BEI Resources while the rest were isolated and cultured gut bacteria from a human fecal sample. The following reagent was obtained through BEI Resources, NIAID, NIH as part of the Human Microbiome Project: Hungatella hathewayi, Strain WAL-18680, HM-308; Streptococcus gallolyticus subsp. gallolyticus, Strain TX20005, HM-272; Bacteroides fragilis, Strain 3_1_12, HM-20; Lachnospiraceae sp., Strain 7_1_58FAA, HM-153; Veillonella sp., Strain 6_1_27, HM-49. The following reagent was obtained through the NIH Biodefense and Emerging Infections Research Resources Repository, NIAID, NIH: Escherichia coli, Strain B6914-MS1, NR-6. DNA from individual cultures were extracted using Qiagen UltraClean kits. The total 16S rRNA gene was amplified (8-27F, 1512-1429R) and sequenced using Sanger sequencing (EtonBio). Clonal sequences were determined through visual inspection of the resultant chromatograms. A combined library of equal amounts of DNA from each isolate was created based on quantifying DNA concentration using Quant-iT dsDNA Assay Kit (Thermo Fisher Scientific). As Quant-iT quantifies total DNA, not just 16S rRNA, qPCR was also used to estimate the resulting mock community composition based on amplifying 16S rRNA. qPCR was performed as follows: the V4 region of the 16S rRNA gene was barcoded and amplified (F515/R806) [19]; all reactions began with a denaturing step of 95C for 2 minutes, followed by 50C for 10 min, followed by 35 amplification cycles - one amplification cycle consists of: 95C for 15 seconds, 60C for 1 minute. PCR was performed using the same primers as qPCR. PCR steps were adapted from Caporaso et al. to permit a variable number of PCR cycles: all reactions began with a denaturing step of 94C for 3 minutes, followed by a variable number of amplification cycles, and finished with 10 minutes of 72C. One amplification cycle consists of: 94C for 45 seconds, 50C for 1 minute, 72C for 1.5 minutes. The number of amplification cycles ranged from 3 to 35. All PCRs were run on a single machine. The order in which the PCRs were done was randomized to avoid systematic bias. 16S rRNA amplicon sequencing was performed using an Illumina MiniSeq with paired-end 150 bp reads.

### Real Community Data Collection

To characterize PCR bias for real microbial communities we analyzed samples from an artificial gut system. Four fecal samples from four separate donors were used to inoculate artificial gut vessels as previously reported [8]. After inoculation, samples from the system were also taken from Day 1, Day 2, and Day 3. Bacterial DNA was extracted using Qiagen DNeasy PowerSoil Kit. The bacterial DNA concentration of the samples was quantified using a Quant-iT dsDNA Assay Kit (Thermo Fisher Scientific). As in the mock community, the V4 region of the 16S rRNA gene was barcoded and amplified. PCR was performed using the same parameters as for the mock community except PCR amplifications were split between 5 machines. The number of amplification cycles ranged from 20 to 35. 16S rRNA amplicon sequencing was done by an Illumina MiniSeq with paired-end 150 bp reads (CAPARASO). After initial data analysis it was found that PCR machine 3 was miscalibrated and the middle amplification step was set to 58C rather than 50C.

### Data Preprocessing

Sequencing data was processed and denoised using DADA2 [20] following a previously published analysis pipeline [8]. For both the mock and real community data, only samples with more than 5000 reads were retained for analysis. This retained 99.7% of sequence variant counts from the mock and 99.8% of sequence variant counts from the real communities respectively. The mock community data was analyzed at the sequence-variant level. Five sequence variants could be uniquely mapped to isolates in the mock community, the other mock community members were amalgamated into a category called “other” for analysis. The real community data was analyzed at the genus level and genera that were not seen in at least 30% of samples with at least 3 counts were amalgamated together into a category called “other” for analysis. Notably, no pseudo-counts were added to the data prior to analysis as the Bayesian multinomial-logistic normal linear model in Equation (5)–(9) models zeros directly [21].

### Analysis of Mock Community Data

To model the mock community data we took *X_i_* (the covariate vector assigned to sample i to be *X_i_* = [1, *x_i_*, *I*_Batch_]*^T^* where 1 represents a constant intercept, *x_i_* denotes the number of PCR cycles that sample i went through, and *I*_Batch_ is a binary variable denoting whether that sample was part of the the first (*I*_Batch_ = 0) or second (*I*_Batch_ = 1) batch of PCR reactions. This specification for *X_i_* implies that Λ can be interpreted as

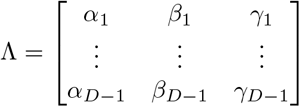

where *α_ℓ_* represents the community composition of the *ℓ*-th log-ratio at cycle 0, and *β_ℓ_* the bias for the *ℓ*-th log-ratio, and *γ_ℓ_* is a variable we introduce to model the effect of batch on the *ℓ*-th log-ratio.

Based on prior reports [4] we choose Bayesian priors that reflected that PCR bias was likely small and centered about zero (no bias) for all log-ratios. This was encoded as Γ = 2*I*_3_ where *I*_3_ represents a 3 × 3 identify matrix and Θ = 0_(*D*−1)×3_. Additionally, our prior reflected our weak belief that the covariance between the absolute abundance of taxa was independent on the log-scale (Ξ = Ψ*I*Ψ*^T^* and *v* = *D* + 2). The multinomial logistic-normal linear model was fit in additive log-ratio coordinates as is default in *stray* and the resulting posterior samples were then transformed into the centered log-ratio coordinate system for figure generation. This transformation was performed using the function *to_clr* provided by the *stray* software package.

### Analysis of Real Community Data

To model the real community data we took *X_i_* to be *X_i_* = [*I*_*P*_1__, …, *I*_*P*_4__, *x_i_*, *I*_PCR_2__, …, *I*_PCR_5__]*^T^* where *I*_*P*_1__ is a binary variable denoting if the *i*-th sample was from person 1, *x*_i_ denotes the PCR cycle number as in the mock community, and *I*_*PCR*_2__ is a binary variable denoting if the i-th sample was amplified on PCR machine number 2.

Based on our analysis of the mock community data we updated our prior to better reflect our updated beliefs. We choose Γ = diag(4, 4, 4, 4, 1, 1, 1, 1, 1) reflecting our updated prior belief regarding the relative scale of the community intercept and other covariates. In this way we used a form of Bayesian sequential learning to update our prior beliefs for the real community data based on the posterior estimates from the mock community analysis. As before we took Θ to be a matrix of zeros reflecting our prior belief that we expect the effect of PCR bias and PCR machine to be small. Ξ and *v* were chosen as in the mock community analysis. The multinomial model was fit and posteriors transformed as in the analysis of the mock community data.

### Predicting Bias from 16S rRNA Amplicon Sequence

A linear regression model predicting PCR bias (*β*) in log-ratio space based on amplicon GC content was fit to 2000 posterior samples. The predictive potential of this model was summarized based on the *R*^2^ statistics for each posterior sample (2.0%, 1.3%–2.6% mean and 25-th–75-th quantiles respectively).

Sequence dissimilarity was summarized as a distance matrix *D*^seq^ with elements 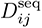 denoting the Levenshtein distance between sequences from taxon *i* and *j* (calculated using function *stringDist* from the R package *Biostrings* [22]). To assess the predictive potential of linear models based on sequence similarity we fit a linear regression model predicting PCR bias in log-ratio space based on the first 10 eigenvectors of the matrix *D*^seq^. The predictive potential of this model was summarized based on the *R*^2^ statistics for each posterior sample (1.6%, 1.5%–1.8% mean and 25-th–75-th quantiles respectively).

To assess the predictive potential of non-linear Gaussian process models we transformed the distance matrix *D*^seq^ into a covariance matrix Σ between sequences using the following radial basis function kernel

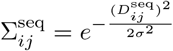

where we refer to *σ* as the bandwidth of the kernel. A similar RBF kernel was built to calculate Σ^GC^ which represents the covariance between taxa based on the squared difference between GC content for two taxa. We assessed the predictive potential of both Σ^seq^ and Σ^GC^ by fitting the following kernel selection model to each posterior sample of bias (*β*^(*s*)^)

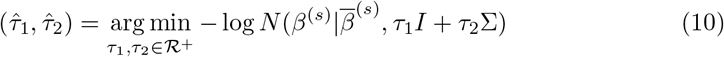

where 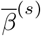 denotes the mean of the biases in posterior sample *s*. We implemented this kernel selection model as the stand-alone R package *NVC* (Normal Variance Components) [23]. To assess the percent of variation explained by either sequence similarity or GC content we defined the following statistic *ρ* = *τ*_2_/(*τ*_1_ + *τ*_2_) which represents the percent of variation explained by the corresponding covariance matrix. Kernel bandwidths were chosen by 10-fold cross validation. In both cases we find that *ρ* ≈ 0.5 with corresponding bandwidths such that Σ ≈ *I* demonstrating that Gaussian processes regression with RBF kernels could not predict PCR bias based on either GC content or sequence similarity.

## Data and Code Availability

Demultiplexed sequencing data was uploaded to SRA (BioProject PRJNA528810 and PRJNA528820). Code to reproduce our analyses along with sequence variant tables from DADA2 are available at https://github.com/jsilve24/pcrbias_paper_code.

## Supporting information

**S1 Fig. Posterior Predictive Checks for Mock Community Analysis** Observed sequence count data *Y* was vectorized and plotted in decreasing order of counts (green line). 2000 samples from the posterior distribution of the multinomial logistic-normal linear model were used to generate 2000 new sequence count datasets. For each observed sequence count, the mean and 95% probability of the corresponding generated datasets is overlayed.

**S2 Fig. Bias Visualized for Mock Community Data** Bias visualized for mock community data. Letting 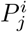 represent the relative abundance of taxon *i* after *j* cycles of PCR we calculated 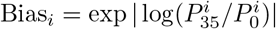. That is, for this visualization we consider a taxon that is decreased by a factor of 1/*b* to have the same bias a taxon increased by a factor of *b*. Here we plot 100 posterior samples of Bias (black lines) and the corresponding mean (green line).

**S3 Fig. Posterior Predictive Checks for Real Community Analysis** Observed sequence count data *Y* was vectorized and plotted in decreasing order of counts (green line). 2000 samples from the posterior distribution of the multinomial logistic-normal linear model were used to generate 2000 new sequence count datasets. For each observed sequence count, the mean and 95% probability of the corresponding generated datasets is overlayed.

**S4 Fig. Bias Visualized for Real Community Data** Bias visualized for mock community data. Letting 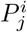 represent the relative abundance of taxon *i* after *j* cycles of PCR we calculated 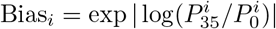. That is, for this visualization we consider a taxon that is decreased by a factor of 1/*b* to have the same bias a taxon increased by a factor of *b*. Here we plot 100 posterior samples of Bias (black lines) and the corresponding mean (green line).

**S5 Fig. Posterior Euclidean Norm of Random Intercept Vector Associated with Each PCR Machine from Real Data Analysis** This norm is shown as a Kernel Density estimate over 2000 posterior samples for each PCR machine.

**S1 File. Marginal Fits for Mulitnomial Logistic-Normal Linear Model Applied to Real Data Analysis** Mean (blue line) and 95% credible regions (grey ribbon) from multinomial logistic-normal linear model (Λ*X*; *Methods*) applied to mock community calibration data. The inferred linear relationship between PCR cycle number and composition is shown adjusting for PCR machine. This adjustment leads the linear fit to appear non-linear even though the overall model is linear. A pseudo-count of 0.65 was added to observed count data prior to log-ratio transformation to enable the data to be visualized along with the posterior estimates. Each column corresponds to one of the four microbial communities analyzed.

## Acknowledgments

We thank Rachel Silverman for her manuscript comments. JDS was supported in part by the Duke University Medical Scientist Training Program (GM007171). JDS and LAD were supported in part by the Global Probiotics Council, a Searle Scholars Award, the Hartwell Foundation, an Alfred P. Sloan Research Fellowship, the Translational Research Institute through Cooperative Agreement NNX16AO69A, the Damon Runyon Cancer Research Foundation, the Hartwell Foundation, and NIH 1R01DK116187-01. SM would like to acknowledge the support of grants NSF IIS-1546331, NSF DMS-1418261, NSF IIS-1320357, NSF DMS-1045153, and NSF DMS1613261.

## References

1. Acinas SG, Sarma-Rupavtarm R, Klepac-Ceraj V, Polz MF. PCR-induced sequence artifacts and bias: insights from comparison of two 16S rRNA clone libraries constructed from the same sample. Applied and Environmental Microbiology. 2005;71(12):8966–8969.

2. Eisenstein M. Microbiology: making the best of PCR bias. Nature Methods. 2018;15:317–320.

3. Love MI, Hogenesch JB, Irizarry RA. Modeling of RNA-seq fragment sequence bias reduces systematic errors in transcript abundance estimation. Nature Biotechnology. 2016;34(12):1287.

4. Polz MF, Cavanaugh CM. Bias in template-to-product ratios in multitemplate PCR. Applied and environmental Microbiology. 1998;64(10):3724–3730.

5. Parada AE, Needham DM, Fuhrman JA. Every base matters: assessing small subunit rRNA primers for marine microbiomes with mock communities, time series and global field samples. Environmental microbiology. 2016;18(5):1403–1414.

6. Gohl DM, Vangay P, Garbe J, MacLean A, Hauge A, Becker A, et al. Systematic improvement of amplicon marker gene methods for increased accuracy in microbiome studies. Nature Biotechnology. 2016;34(9):942.

7. Silverman JD, Roche K, Holmes ZC, David LA, Mukherjee S. Bayesian Multinomial Logistic Normal Models through Marginally Latent Matrix-T Processes. arXiv e-prints. 2019; p. arXiv:1903.11695.

8. Silverman JD, Durand HK, Bloom RJ, Mukherjee S, David LA. Dynamic linear models guide design and analysis of microbiota studies within artificial human guts. Microbiome. 2018;6(1):202. doi:10.1186/s40168-018-0584-3.

9. Gloor GB, Wu JR, Pawlowsky-Glahn V, Egozcue JJ. It’s all relative: analyzing microbiome data as compositions. Annals of Epidemiology. 2016;26(5):322–329.

10. Silverman JD, Washburne AD, Mukherjee S, David LA. A phylogenetic transform enhances analysis of compositional microbiota data. eLife. 2017;6. doi:10.7554/eLife.21887.

11. Love JL, Scholes P, Gilpin B, Savill M, Lin S, Samuel L. Evaluation of uncertainty in quantitative real-time PCR. Journal of microbiological methods. 2006;67(2):349–356.

12. Kanagawa T. Bias and artifacts in multitemplate polymerase chain reactions (PCR). Journal of bioscience and bioengineering. 2003;96(4):317–323.

13. Browne HP, Forster SC, Anonye BO, Kumar N, Neville BA, Stares MD, et al. Culturing of ‘unculturable’human microbiota reveals novel taxa and extensive sporulation. Nature. 2016;533(7604):543.

14. McLaren MR, Willis AD, Callahan BJ. Consistent and correctable bias in metagenomic sequencing measurements. bioRxiv. 2019;doi:10.1101/559831.

15. Illumina. An introduction to next-generation sequencing technology. 2015;.

16. Suzuki MT, Giovannoni SJ. Bias caused by template annealing in the amplification of mixtures of 16S rRNA genes by PCR. Appl Environ Microbiol. 1996;62(2):625–630.

17. Gloor GB, Macklaim JM, Vu M, Fernandes AD. Compositional uncertainty should not be ignored in high-throughput sequencing data analysis. Austrian Journal of Statistics. 2016;45(4):73. doi:10.17713/ajs.v45i4.122.

18. Silverman JD. stray: Multinomial Logistic Normal Linear Models. GitHub. 2019;.

19. Caporaso JG, Lauber CL, Walters WA, Berg-Lyons D, Lozupone CA, Turnbaugh PJ, et al. Global patterns of 16S rRNA diversity at a depth of millions of sequences per sample. Proceedings of the national academy of sciences. 2011;108(Supplement 1):4516–4522.

20. Callahan BJ, McMurdie PJ, Rosen MJ, Han AW, Johnson AJ, Holmes SP. DADA2: High-resolution sample inference from Illumina amplicon data. Nature Methods. 2016;13(7):581–3. doi:10.1038/nmeth.3869.

21. Silverman JD, Roche K, Mukherjee S, David LA. Naught all zeros in sequence count data are the same. bioRxiv. 2018;doi:10.1101/477794.

22. Pages H, Aboyoun P, Gentleman R, DebRoy S. Biostrings: Efficient manipulation of biological strings. R package version. 2017;2(0).

23. Silverman JD. NVC: Normal Variance Components. GitHub. 2019;.

