## Supplemental Figures for "Measuring and Mitigating PCR Bias in Microbiome Data"

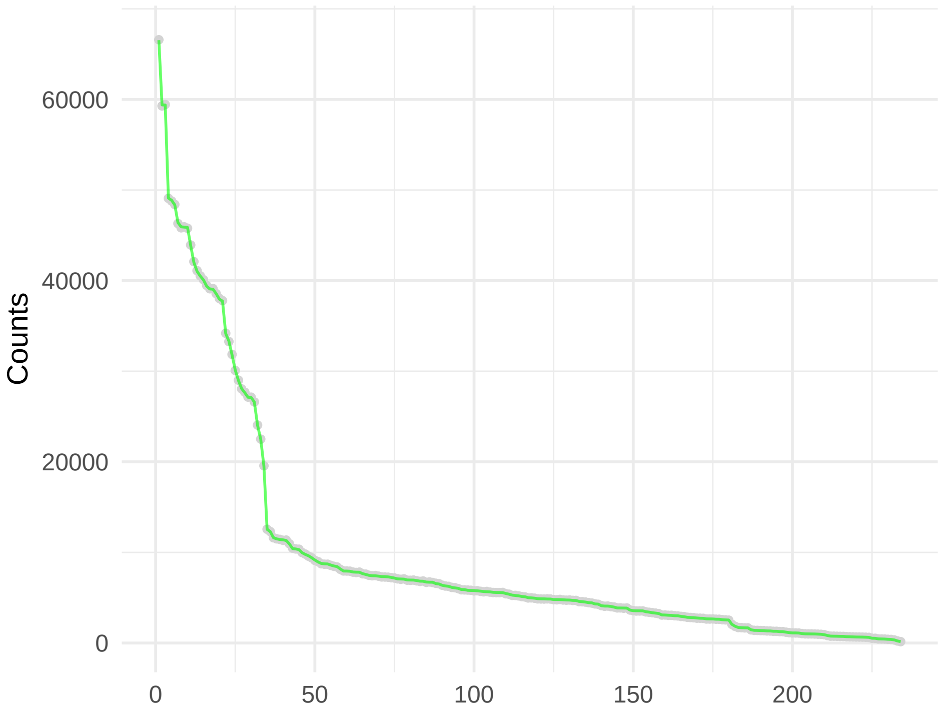

Figure S1: Posterior Predictive Checks for Mock Community Analysis

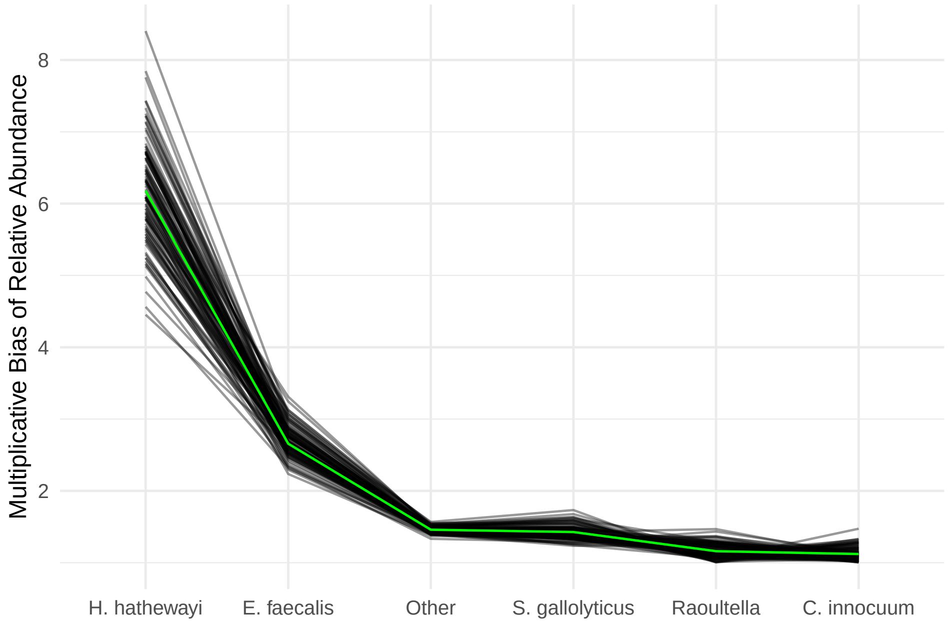

Figure S2: Bias Visualized for Mock Community Data

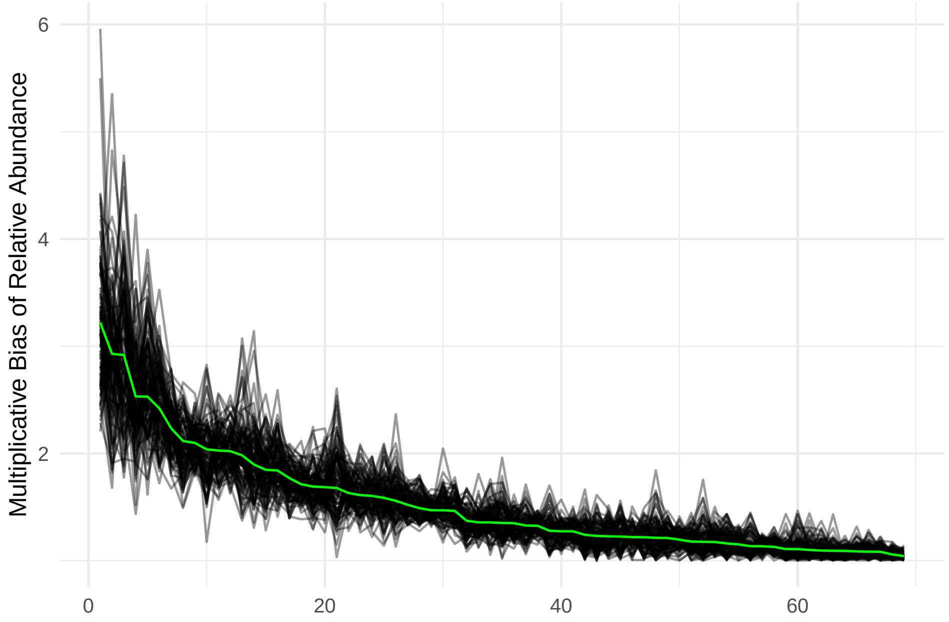

Figure S3: Bias Visualized for Real Community Data

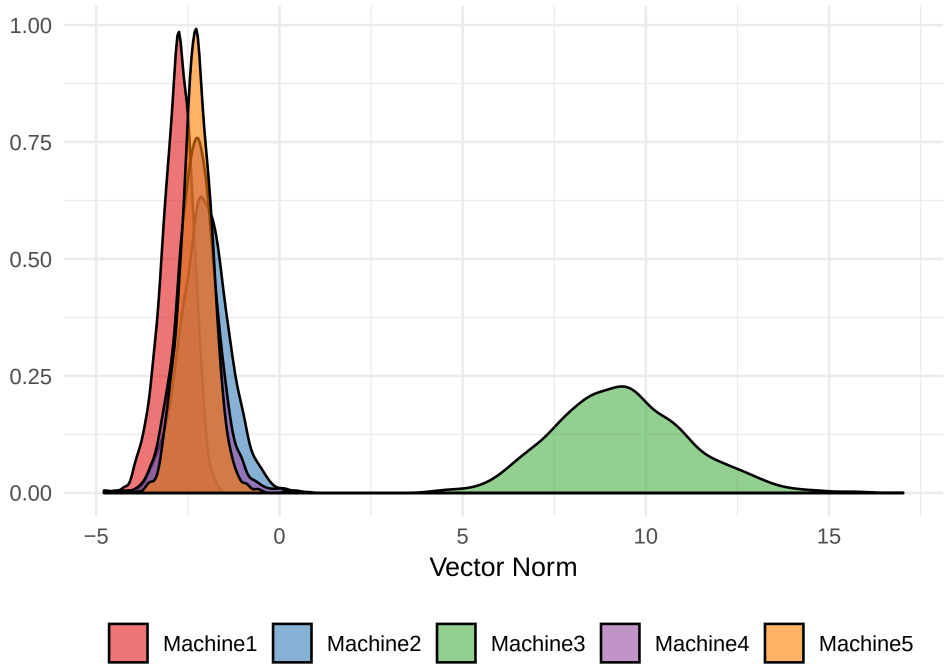

Figure S4: Posterior Euclidean Norm of Random Intercept Vector Associated with Each PCR Machine from Real Data Analysis

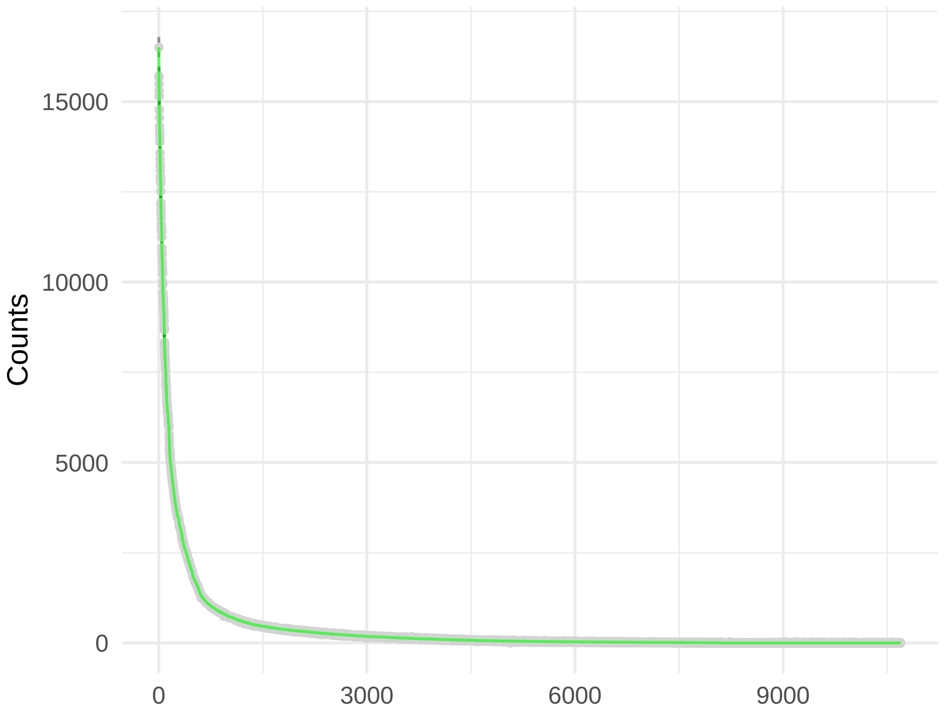

Figure S5: Posterior Predictive Checks for Real Community Analysis
