## Supplementary figures and images for "Measuring and Mitigating PCR Bias in Microbiome Data"

### Supplemental File 1

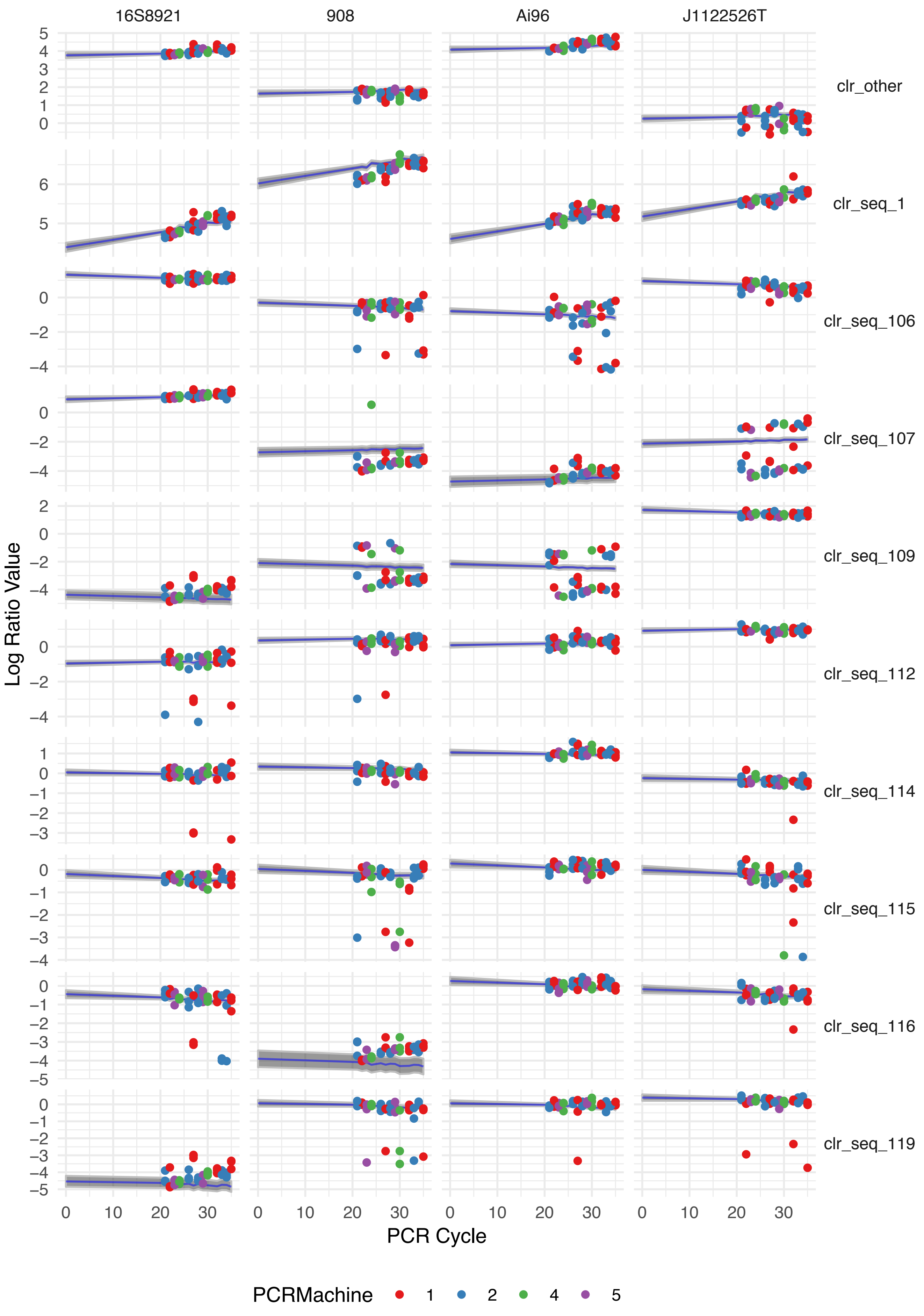

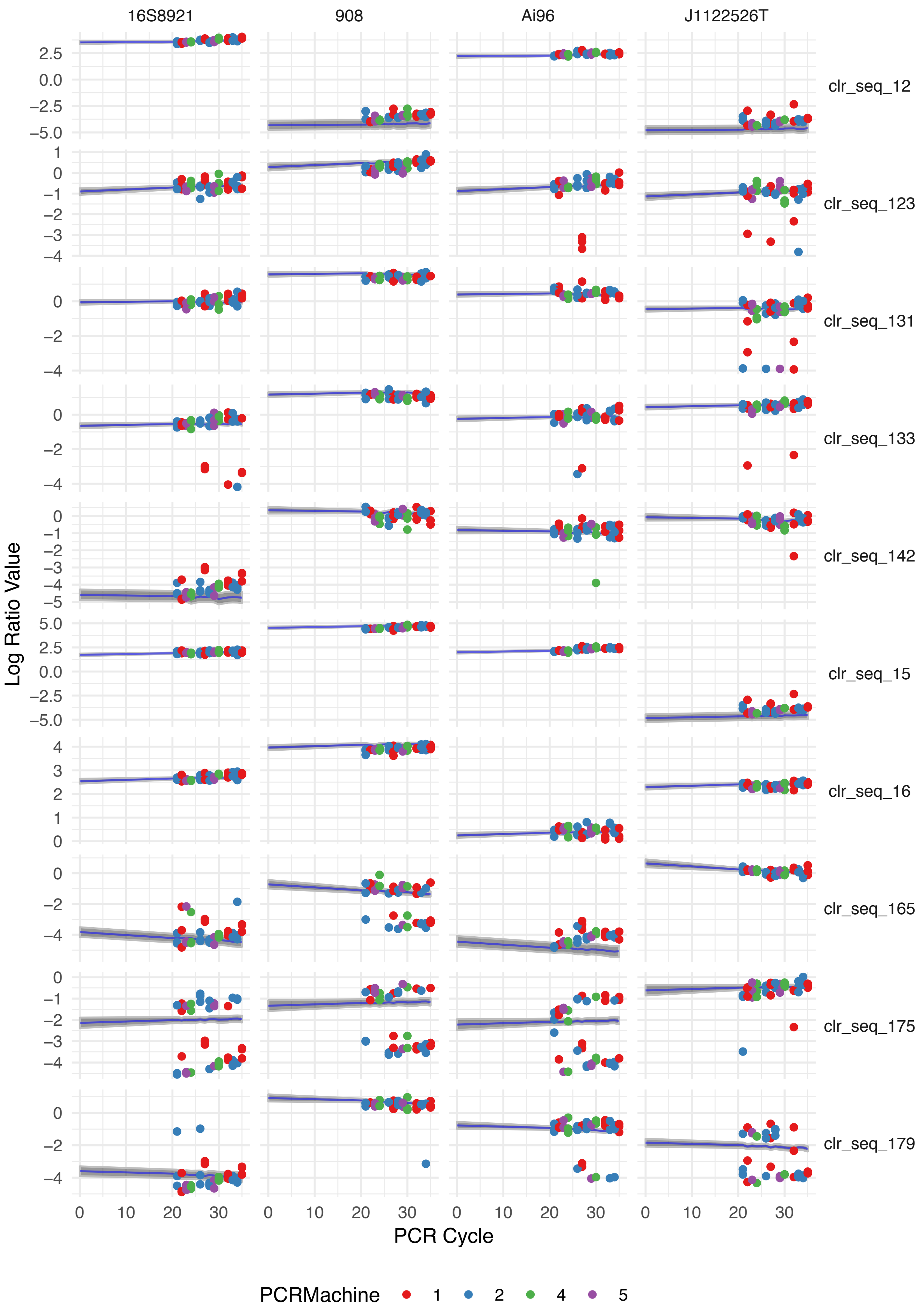

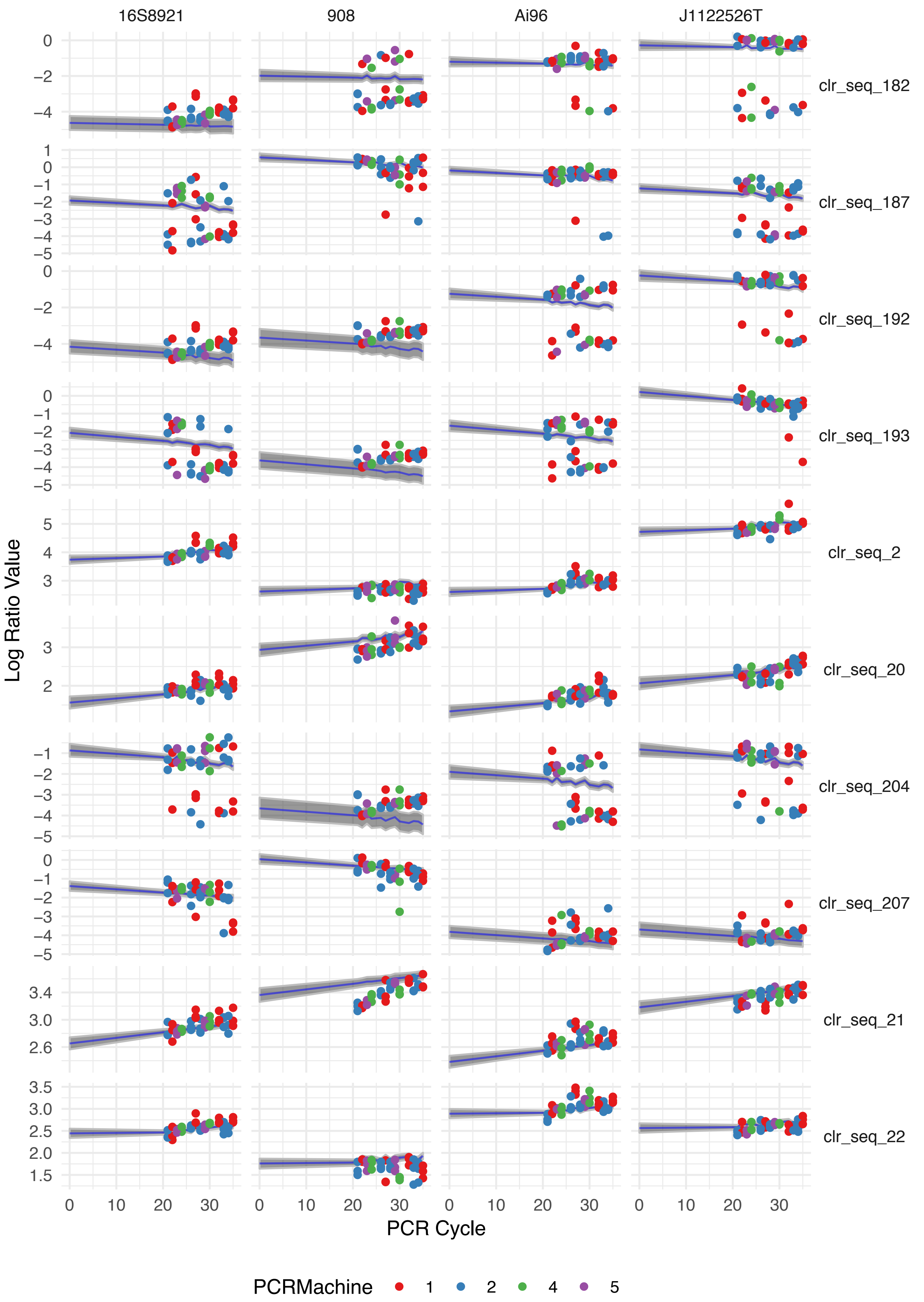

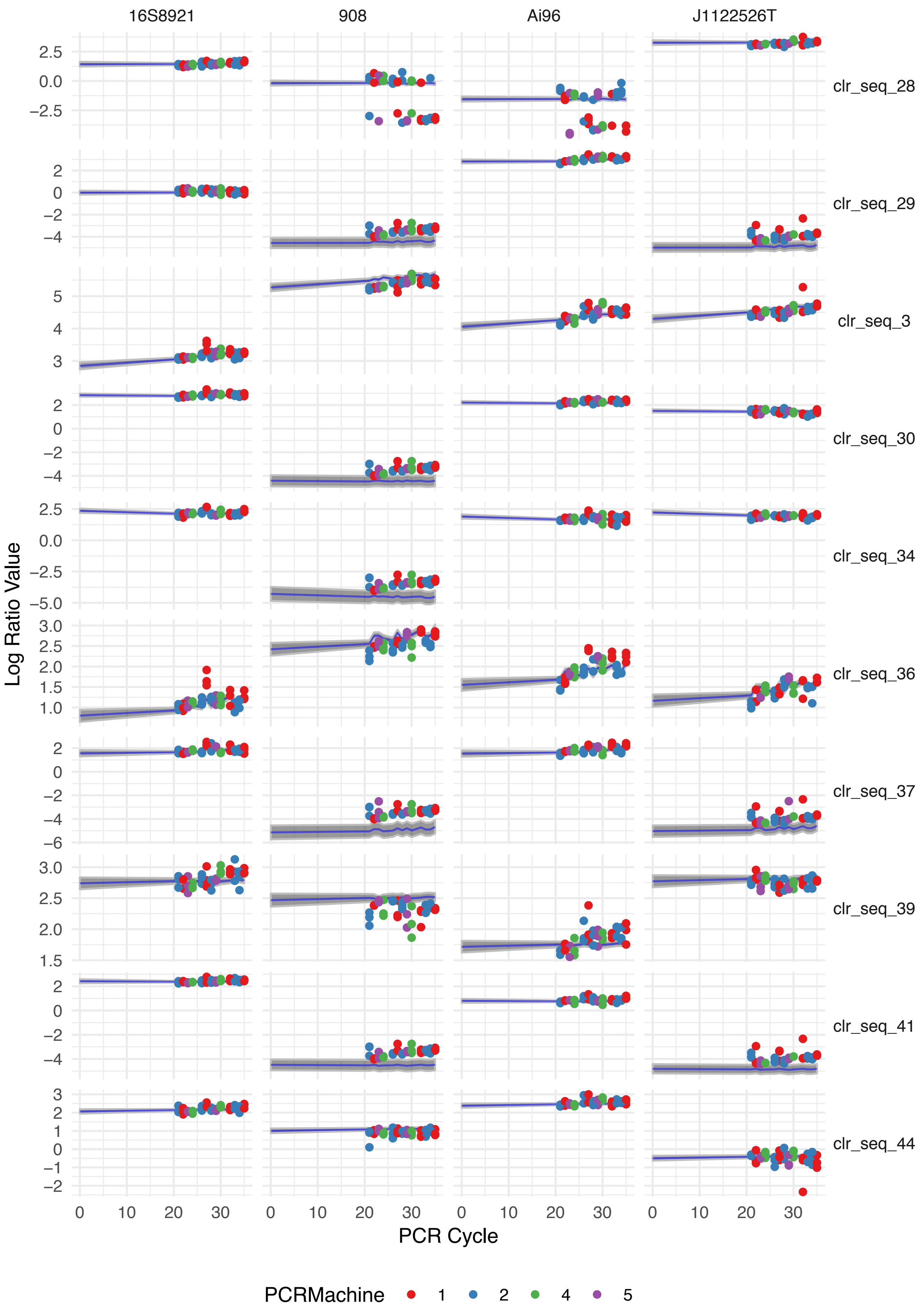

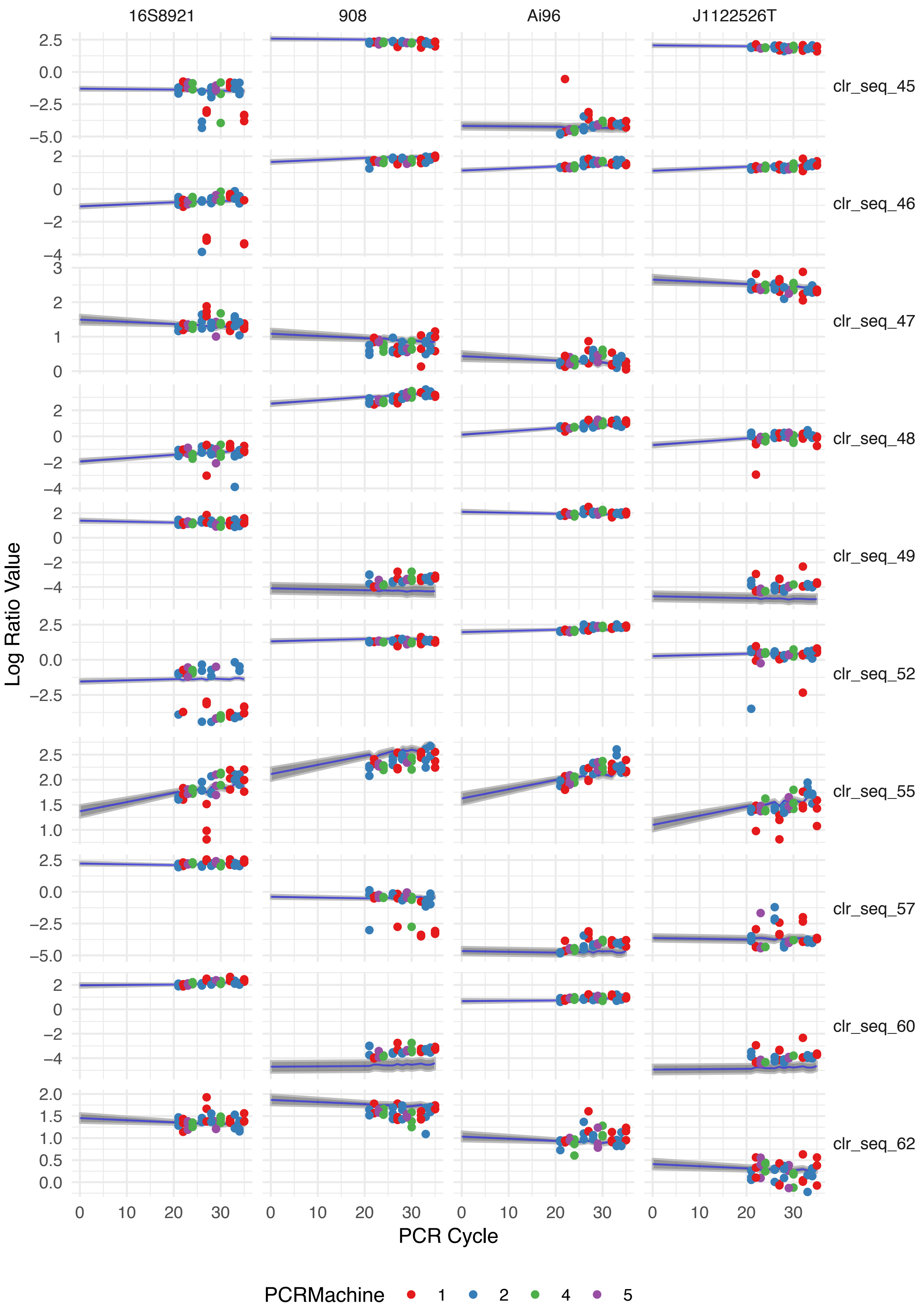

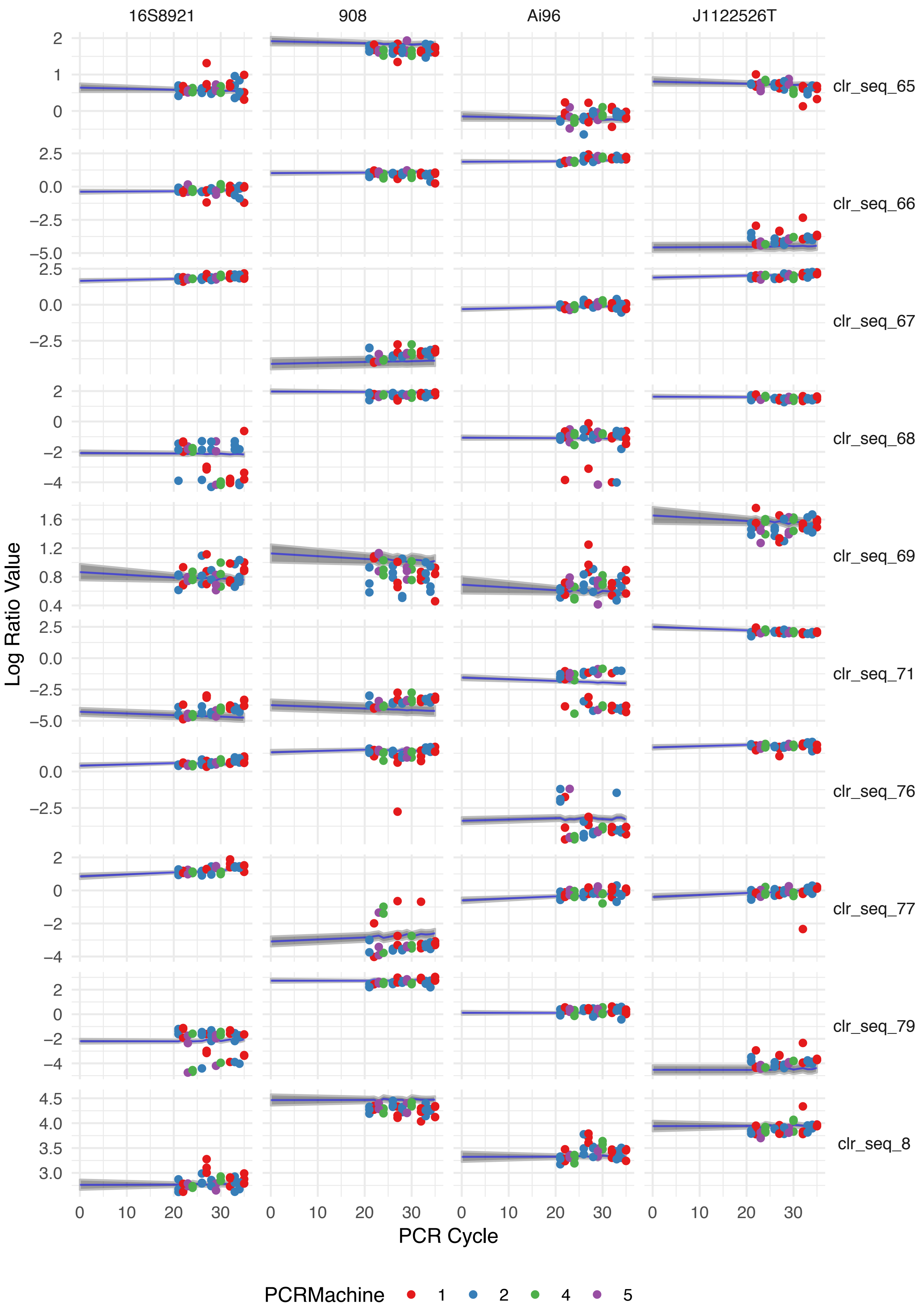

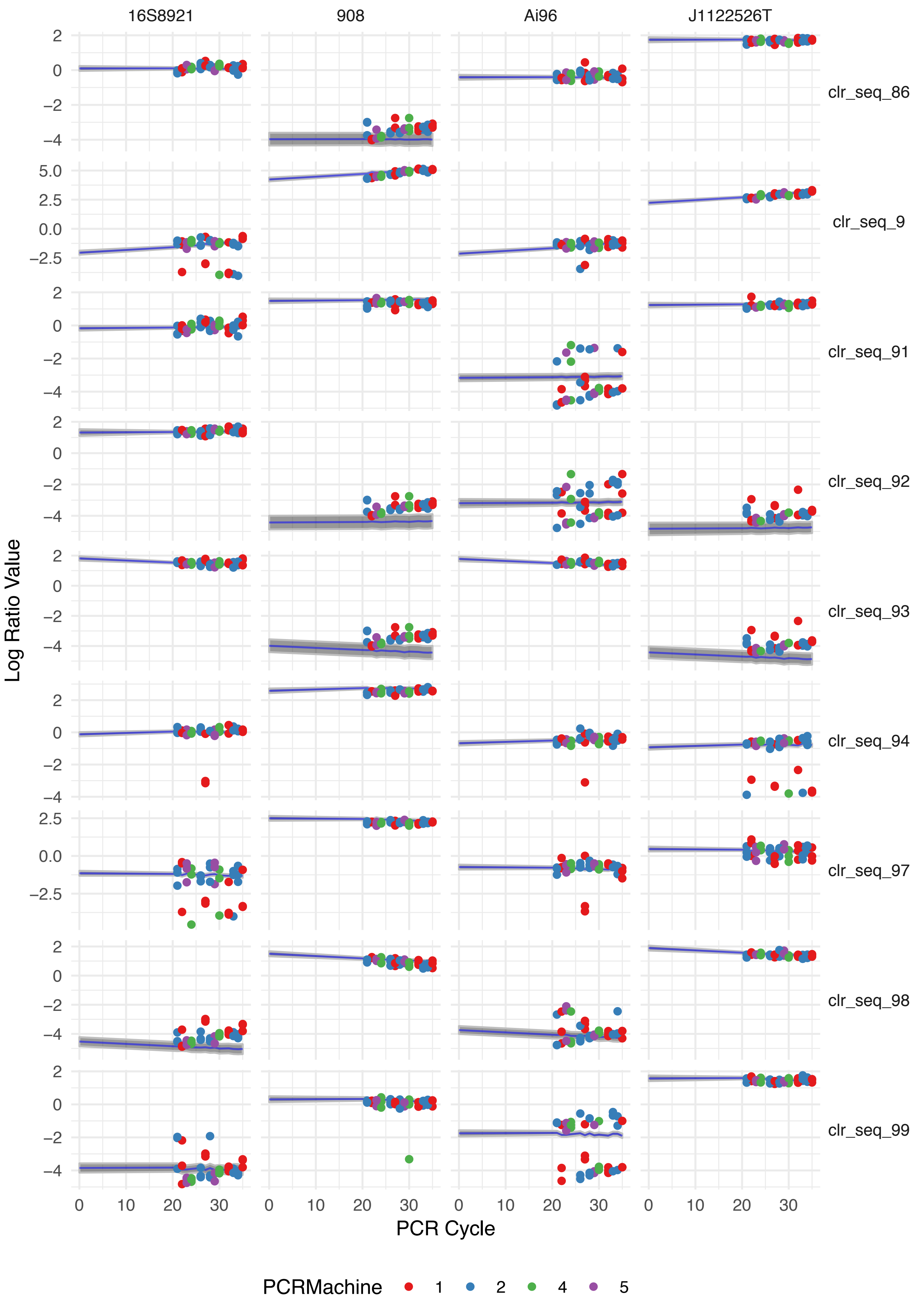
